# Distinguishing Specific from Broad Genetic Associations between External Correlates and Common Factors

**DOI:** 10.1101/2025.06.02.657362

**Authors:** Javier de la Fuente, Diego Londoño-Correa, Elliot M. Tucker-Drob

## Abstract

Within the Genomic SEM framework, common factors are often used to index shared genetic etiology across constellations of GWAS phenotypes. A standard common pathway model, in which a genetic association is estimated between an external GWAS phenotype and a common factor, assumes that all genetic associations between the external GWAS phenotype and the individual indicator phenotypes are mediated through the factor. This assumption can be tested using the QTrait statistic, which compares the common pathway model to an independent pathways model that allows for direct genetic associations between the external GWAS phenotype and the individual indicators of the factor. We expand upon the QTrait approach by describing an effect size index that quantifies the degree to which the common pathways model is violated, and we provide a systematic approach for empirically identifying specific direct pathways between an external trait and indicator traits. Our method comprises a series of omnibus tests and outlier detection algorithms indexing the heterogeneity of associations between the genetic component of external traits and the individual indicators of common factors. We provide a set of automated functions which we apply to investigate the patterns of genetic associations across a set of external correlates with respect to indicators of general cognitive ability and case-control and proxy GWAS indices of Alzheimer’s disease.

## Introduction

Recent multivariate genome-wide associations studies (GWASs) have revealed shared genetic architecture across diverse constellations of biobehavioral traits. Within the Genomic SEM framework (Grotzinger et al., 2019), common factors are often used to index and characterize shared genetic architecture at different levels of analysis. Common factors account for shard genetic etiology across multiple GWAS traits, providing a parsimonious representation of the empirical patterns of genetic covariance across the traits. In many applications, a key goal is to estimate genetic relations between these common factors and external GWAS traits. This is especially appealing because it reduces the number of genetic correlations (or genetic regression coefficients) estimated, and consolidates signal (and thus power) across indicators of a given factor (Grotzinger et al., 2022; Schwaba et al., 2025).

However, a key assumption of such a model is that the genetic associations between the external trait and the individual indicators is well-represented by associations between the external trait and the factor(s) on which the indicators load. Put differently, the standard common pathway model assumes that all genetic associations between the external GWAS correlates and the individual indicator phenotypes are mediated through the common factor. In instances in which the model fails to adequately capture the underlying pattern of associations, additional associations between the external trait and one or more individual indicators (direct effects) may be warranted. Grotzinger et al. (2022) introduced the QTrait statistic as an omnibus test of heterogeneity of relations between an external trait and indicator traits beyond that implied by the common pathway model. QTrait is a χ^2^ distributed test statistic that is estimated by comparing a model with pathways from the external GWAS correlate to the factor with a model in which the external correlate has direct pathways to the individual indicator phenotypes that load onto the common factor. A significant QTrait statistic indicates that the relationship between the GWAS indicator traits and the external correlate cannot be solely explained by pathways through the common factor(s), implying the existence of more specific underlying pathways. If such specific pathways are omitted from the model, the estimate of the association between the trait and the factor may be biased.

The QTrait statistic can detect heterogeneous associations between common factors and external correlates, but it is not designed to identify either the source or the magnitude of this heterogeneity. As an omnibus inferential test, it evaluates heterogeneity based on statistical significance, without accounting for effect size. Ignoring effect size may lead to the detection of trivial differences in high-powered GWAS studies, while in low-powered studies, meaningful differences might be mistaken for sampling variation. Furthermore, identifying outliers can provide valuable insights into specific GWAS phenotypes that follow distinct pathways of association with external correlates. This identification is crucial, as it may reveal unique genetic pathways linking individual traits to external correlates, providing a foundation for generating testable hypotheses about the genetic mechanisms underlying complex traits.

Here, we provide a systematic approach to investigate and correct for potential heterogeneity of associations between the genetic component of external correlates and the individual indicators of common factors of shared genetic architecture. Our approach combines both statistical significance and effect size measures of heterogeneity (lSRMR), and provides a systematic approach for identifying the specific indicator phenotypes for which direct pathways may be added to the standard common pathway model. Moreover, we provide a set of automated functions within the Genomic SEM framework and illustrate their application to investigate the patterns of associations across a set of external correlates with respect to general cognitive function and Alzheimer’s disease (AD). We used the QTrait function to explore the patterns of genetic associations across a set of external biobehavioral correlates with respect to two genetic factors representing: 1) genetic propensity towards AD using case-control GWAS of AD and proxy-phenotype GWAX of maternal and paternal history of AD, and 2) a genetic general intelligence factor (*g*) indexing shared genetic variation among seven cognitive traits. For AD we used the Genomic SEM-based multivariate model proposed by de la Fuente et al. (2022) to represent the overall genetic predisposition towards AD as a common factor influencing both the direct GWAS phenotype and two GWAX phenotypes. For general cognitive function we used the genetic general intelligence factor (*g*) described in de la Fuente et al. (2021).

## Results

### Overview of the QTrait function

The QTrait function uses the QTrait statistic introduced by Grotzinger et al. (2022) as an omnibus statistical test of heterogeneity. The QTrait is based on χ^2^ difference tests comparing two competing models: 1) the common pathway model, where the external correlate predicts only the common factor, and 2) the independent pathways model, where the external correlate predicts the individual indicator phenotypes (**Figure 1**). The χ^2^ difference between these two models is referred to as the QTrait statistic. A significant QTrait statistic suggests that the relationship between the individual indicator phenotypes and the external correlate cannot be solely explained by the common factor, indicating more specific pathways to individual indicator phenotypes. To determine statistical significance for the QTrait statistic we used a Bonferroni-corrected *p*-value threshold based on the number of external correlates for which the function independently computes the heterogeneity indices.

**Figure 1.**
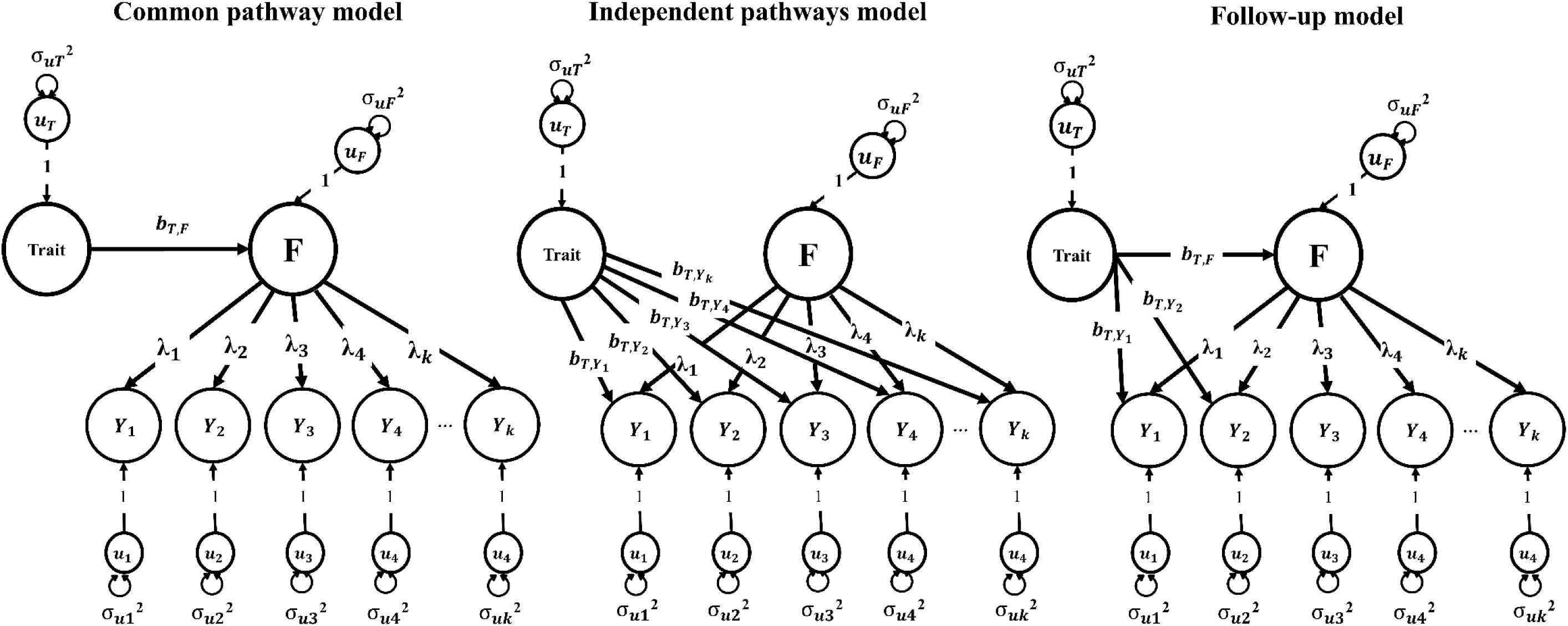
The fit of the common pathway and independent pathways models are compared for producing the QTrait heterogeneity index. The follow-up model is automatically re-specified by the QTrait function, allowing unconstrained direct paths to the identified outlier traits (in this case Y1 and Y2)

In addition to the QTrait heterogeneity index, the QTrait function computes the local standardized root mean squared residual (lSRMR), a global effect size index of heterogeneity which quantifies the magnitude of the discrepancies between the empirical Linkage Disequilibrium Score Regression (LDSC)-derived genetic correlations and those implied by the common pathway model (see Methods). We considered localSRMR values equal to or exceeding 25% of the of the root mean square genetic correlation between the external correlate and the individual indicator phenotypes to reflect non-negligible differences between the observed and the model-implied genetic correlations, thus reflecting meaningful heterogeneity in the association between the factor and the external correlate.

We applied a three-criteria test to assess heterogeneity in the associations between the external correlate and the individual indicator phenotypes. Specifically, we required that: (1) the genetic correlation between the external correlate and the common factor surpasses a Bonferroni-corrected *p*-value threshold; (2) the QTrait heterogeneity index is statistically significant at the same threshold; and (3) the lSRMR effect size heterogeneity index exceeds predefined thresholds, with the default criteria being lSRMR > 0.10 and greater than 25% of the root mean square genetic correlation between the external correlate and the indicator phenotypes that load on the factor.

For external correlates showing heterogeneous associations with the common factor, as indicated by the three-criteria test, we identified specific outlier phenotypes that deviated from the common pathway model. An indicator phenotype was considered an outlier if the residual genetic correlation with the external correlate exceeded 0.10 and was greater than 25% of the root mean square genetic correlation across all indicator phenotypes and the external correlate. When outliers were detected, we fitted a series of follow-up models with unconstrained direct paths from the external correlate to the most extreme outlier, iterating until the model was saturated (i.e., df = 0) or until no further significant heterogeneity was detected.

### Alzheimer’s Disease

In the analysis, 14 external correlates were considered, out of which five exhibited significant associations with the Alzheimer’s Disease (AD) factor at the Bonferroni-corrected *p*-value threshold of *p* < .004 (**Table 1**). These include educational attainment, cognitive performance, high-density lipoprotein (HDL), coronary artery disease (CAD), and heart failure. Among these, HDL (rG=0.11, SE=0.04, *p*=.002), CAD (rG =-0.15, SE=0.04, *p*=.001), and heart failure (rG =-0.19,SE=0.06, *p*=.001) did not exhibit significant deviations from the common pathway model according to the omnibus tests of heterogeneity, suggesting that these external correlates are broadly relevant for both the direct GWAS and the family GWAX AD phenotypes (**Table 1**). On the contrary, we found evidence of heterogeneity in the associations between educational attainment (QTrait (2)=36.75, *p*<.001; lSRMR=.13), cognitive performance (QTrait (2)=26.61, *p*<.001; lSRMR=.11) and the individual AD phenotypes, indicating that these external correlates may operate in more specific pathways with respect to direct GWAS and proxy GWAX of AD. The outlier detection method indicated that maternal GWAX presented significant deviations from the common pathway model (**Figure 2**). More specifically, maternal GWAX of AD presented stronger positive associations with both EA and cognitive performance than those implied by the common pathway model. The follow-up models for educational attainment (QTrait (1)= 5.25, *p*=.022; lSRMR=.08) and cognitive performance (QTrait (1)= 1.57, *p*=.260; lSRMR=.06) with unconstrained direct paths to maternal AD GWAX did not reveal significant residual heterogeneity, while the associations between the AD factor remained statistically significant for both educational attainment (β =-0.16, SE=0.03, *p*=.001) and cognitive performance (β =-0.23,SE=0.04, *p*<.001).

**Figure 2.**
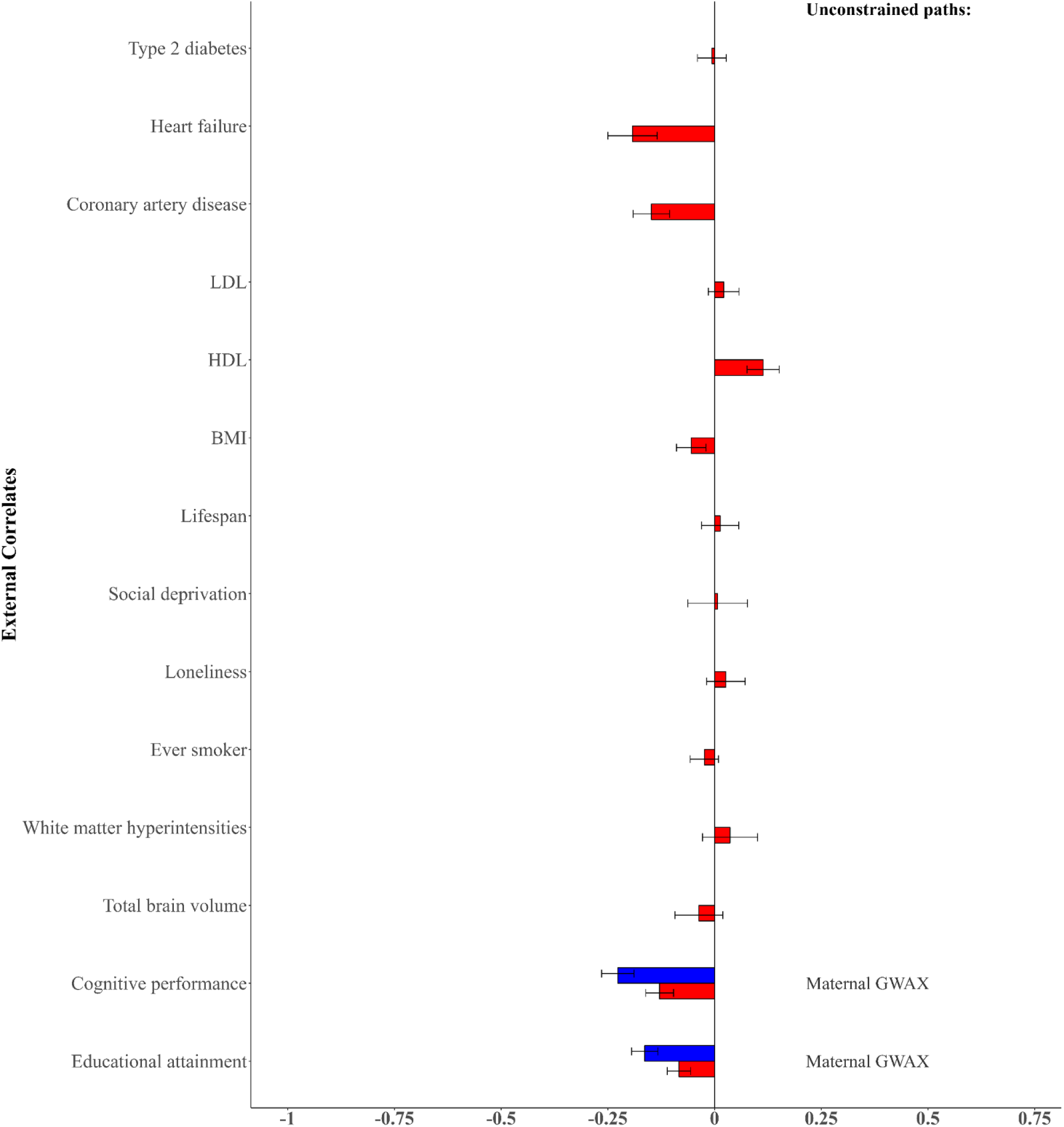
Genetic correlations across external correlates and the Alzheimer’s Disease (AD) factor. The common pathway model assumes that all genetic associations between the external GWAS correlates and the individual indicator phenotypes are mediated through the common factor. The follow-up model corresponds with a model with unconstrained direct pathways to the outlier traits presenting significant deviations from the expectations under the common pathway model according to the three-test criteria of heterogeneity (see Methods).

**Table 1.** Heterogeneity statistics and patterns of associations between case-control GWAS of Alzheimer’s disease (AD) and proxy-phenotype GWAX of maternal and paternal history of AD in relation to 14 biobehavioral external correlates.

| External correlate | Common pathway model |  |  | Follow-up model |  |  | Outliers |
| --- | --- | --- | --- | --- | --- | --- | --- |
|  | rG (SE) | QTrait (df) | ISRMR | rG (SE) | QTrait (df) | ISRMR |  |
| Educational attainment | -0.08* (0.03) | 36.75* (2) | 0.13<sup>a</sup> | -0.16* | 5.25 (1) | 0.08<sup>a</sup> | Maternal GWAX |
| Cognitive performance | -0.13* (0.03) | 26.61* (2) | 0.13<sup>a</sup> | -0.23* | 1.57 (1) | 0.06<sup>a</sup> | Maternal GWAX |
| Total brain volume | -0.04 (0.06) | 2.24 (2) | 0.08<sup>a</sup> | - | - | - | - |
| White matter hyperintensities | 0.04 (0.06) | 0.53 (2) | 0.05<sup>a</sup> | - | - | - | - |
| Ever smoker | -0.02 (0.03) | 4.14 (2) | 0.06<sup>a</sup> | - | - | - | - |
| Loneliness | 0.03 (0.05) | 2.09 (2) | 0.05<sup>a</sup> | - | - | - | - |
| Social deprivation | 0.01 (0.07) | 0.01 (2) | 0.01<sup>a</sup> | - | - | - | - |
| Lifespan | 0.01 (0.04) | 3.49 (2) | 0.09<sup>a</sup> | - | - | - | - |
| BMI | -0.05 (0.03) | 0.23 (2) | 0.01 | - | - | - | - |
| HDL | 0.11* (0.04) | 3.11 (2) | 0.06<sup>a</sup> | - | - | - | - |
| LDL | 0.02 (0.04) | 3.69 (2) | 0.03<sup>a</sup> | - | - | - | - |
| Coronary artery disease | -0.15* (0.04) | 4.01 (2) | 0.09<sup>a</sup> | - | - | - | - |
| Heart failure | -0.19* (0.06) | 0.53 (2) | 0.05 | - | - | - | - |
| Type 2 diabetes | -0.01 (0.03) | 1.90 (2) | 0.06<sup>a</sup> | - | - | - | - |

rG = genetic correlation. *ISRMR* = local standardized mean squared residual.
\*Statistically significant at the Bonferroni-corrected p-value threshold of $p < .004$ .
<sup>a</sup>*ISRMR* surpassing 25% of the root mean square genetic correlation between the external correlate and the individual traits loading on the common factor.

Figure 3 contains plots presenting (A) the observed, model-implied (common pathway model), and residual genetic correlations between educational attainment and the three indicators of the Alzheimer’s disease (AD) liability factor—maternal GWAX, paternal GWAX, and case-control GWAS of AD, and B) a scatterplot of unstandardized beta coefficients of educational attainment on the three indicators against their respective unstandardized factor loadings on the AD liability factor (de la Fuente et al., 2022). The size of each dot corresponds to the inverse of the variance of the unstandardized beta coefficient. The figure also shows the model-implied association between educational attainment and AD liability from both the common pathway and the unconstrained model with direct path to maternal GWAX, represented by the slope of the regression line.

**Figure 3.**
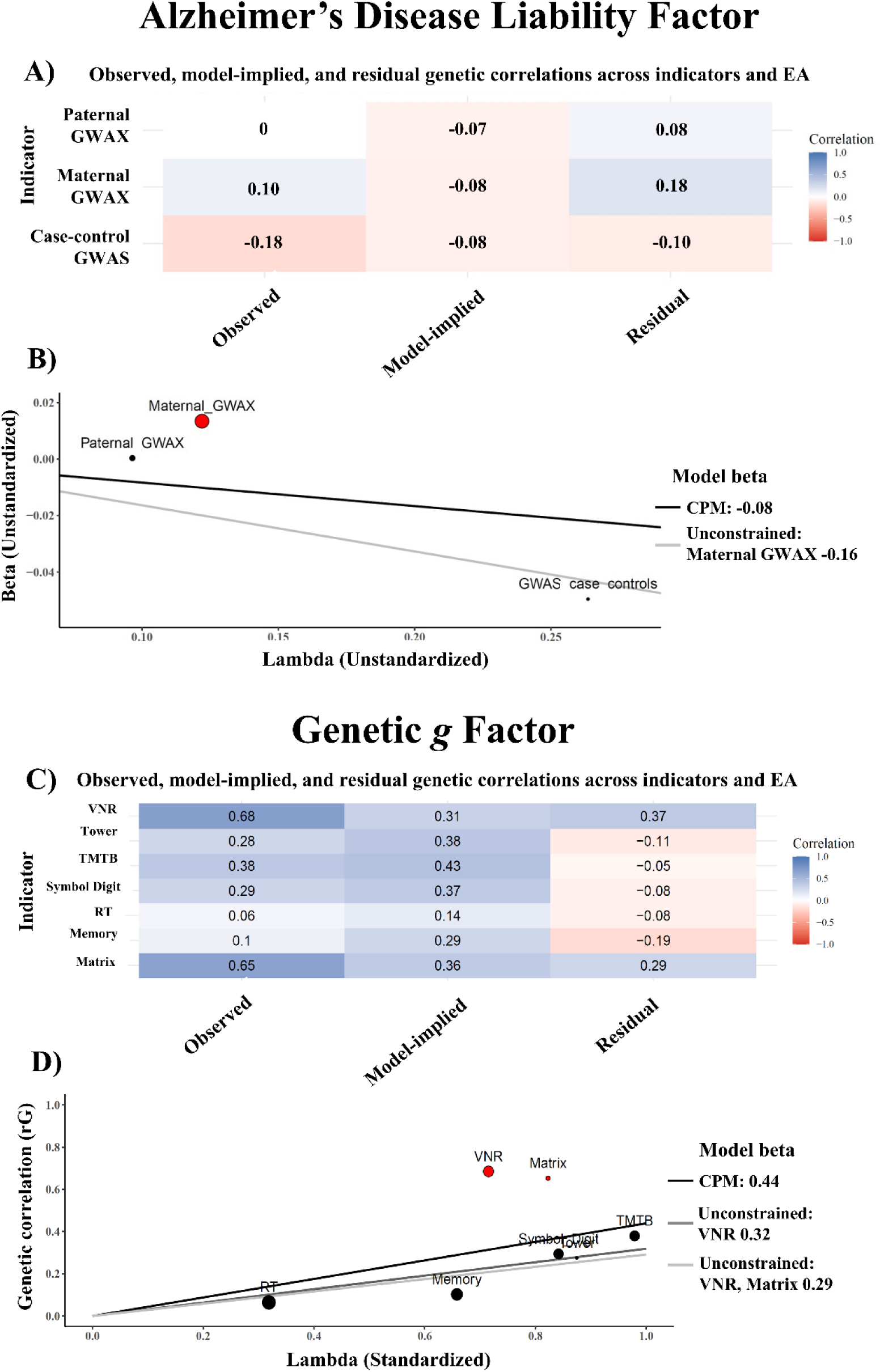
**A**) Observed, model-implied, and residual genetic correlations between the three AD indicators and educational attainment (EA). **B**) Scatterplot of unstandardized betas of EA on AD indicators against their unstandardized loadings on the AD factor (de la Fuente et al., 2021); dot size reflects inverse variance. **C**) Observed, model-implied, and residual genetic correlations between the seven genetic *g* cognitive indicators and EA. **D**) Scatterplot of genetic correlations between EA and the cognitive indicators against their standardized loadings on the genetic *g* factor; dot size reflects inverse variance. CPM = common pathway model.

The common pathway model estimates the genetic association between educational attainment and AD liability but fails to fully account for the relations with maternal GWAX indicator, as indicated in Figure 3 by the deviation of the maternal GWAX from the line representing the model-implied association from the common pathway model, and the large residual genetic correlation between maternal GWAX and educational attainment (0.18). If the common pathway model were sufficient, genetic correlations with educational attainment would scale proportionally with the factor loadings of the indicators on the AD liability factor. However, maternal GWAX deviates from this expectation, indicating stronger-than-expected positive genetic correlation with educational attainment. This suggests that the association between educational attainment and maternal GWAX involves additional genetic effects beyond those captured by the common pathway model. These findings highlight the limitations of the common pathway and imply that the genetic relationships between educational attainment and AD differs depending on whether direct case-control GWAS versus proxy GWAX of parental history are used. It is possible that the maternal GWAX in particular suffers from a pernicious form of ascertainment bias. It can be seen from Figure 3 that when the model is relaxed to account for the specific genetic relation between educational attainment and the maternal GWAX, the model-implied genetic association between educational attainment and AD doubles in strength.

### Cognitive Function

We further investigated the generality versus specificity of associations between 20 external GWAS correlates (the same 14 traits analyzed in AD, and 6 additional neuropsychiatric phenotypes associated with cognitive function) and the genetic general intelligence factor (*g*) described in de la Fuente et al. (2021). The genetic *g* factor accounts for the shared genetic variance among 7 continuous cognitive traits GWAS phenotypes. Out of the 20 external correlates considered, 15 exhibited significant associations with the genetic g factor at the Bonferroni-corrected *p*-value threshold of *p* < .002 (**Table 2**). The 5 external correlates that exhibited significant heterogeneity were: educational attainment (EA), cognitive performance, loneliness, lifespan, and attention-deficit/hyperactivity disorder (ADHD), suggesting more specific pathways with respect to genetic *g*.

**Table 2.** Heterogeneity statistics and patterns of associations between the g factor in relation to 20 biobehavioral external correlates.

| External correlate | Common pathway model |  |  | Follow-up model |  |  | Outliers |
| --- | --- | --- | --- | --- | --- | --- | --- |
|  | rG (SE) | QTrait (6) | ISRMR | rG (SE) | QTrait (df) | ISRMR |  |
| Schizophrenia | -0.39 | 17.40 | 0.06 | - | - | - | - |
| Bipolar disorder | -0.25 | 32.26 | 0.07 | - | - | - | - |
| Autism | 0.08 | 62.21 | 0.10<sup>a</sup> | - | - | - | - |
| ADHD | -0.28 | 202.12 | 0.13<sup>a</sup> | -0.14 | 31.47 (4) | 0.07 | VNR, Matrix |
| Openness to experience | 0.02 | 70.55 | 0.12<sup>a</sup> | - | - | - | - |
| Conscientiousness | -0.12 | 14.48 | 0.08 | - | - | - | - |
| Educational attainment | 0.44 | 1424.50 | 0.20<sup>a</sup> | 0.29 | 40.23 (4) | 0.06 | VNR, Matrix |
| Cognitive performance | 0.82 | 2892.15 | 0.18<sup>a</sup> | 0.72 | 49.18 (5) | 0.13 | VNR |
| Total brain volume | 0.21 | 20.82 | 0.07 | - | - | - | - |
| White matter hyperintensities | -0.11 | 9.02 | 0.05 | - | - | - | - |
| Ever smoker | -0.13 | 73.22 | 0.09 | - | - | - | - |
| Loneliness | -0.15 | 43.83 | 0.07 | - | - | - | - |
| Social deprivation | -0.23 | 64.72 | 0.12<sup>a</sup> | -0.22 | 69.12 (5) | 0.10 | Matrix |
| Lifespan | 0.25 | 51.93 | 0.11<sup>a</sup> | 0.24 | 41.75 (5) | 0.07 | Matrix |
| BMI | -0.07 | 101.61 | 0.10 | - | - | - | - |
| HDL | 0.04 | 23.73 | 0.06 | - | - | - | - |
| LDL | 0.00 | 23.82 | 0.06 | - | - | - | - |
| Coronary artery disease | -0.11 | 42.95 | 0.07 | - | - | - | - |
| Heart failure | -0.16 | 27.01 | 0.09 | - | - | - | - |
| Type 2 diabetes | -0.07 | 58.63 | 0.07 | - | - | - | - |

rG = genetic correlation. *ISRMR* = local standardized mean squared residual.
\*Statistically significant at the Bonferroni-corrected p-value threshold of $p < .004$ .
<sup>a</sup>*ISRMR* surpassing 25% of the root mean square genetic correlation between the external correlate and the individual traits loading on the common factor.

We focused our attention on the 3 external traits presenting the highest genetic correlations with the genetic *g* factor: educational attainment (rG=.44, *p*<.001), cognitive performance (rG=.82, *p*<.001), and ADHD (rG=-.28, *p*<.001). The outlier detection method highlighted verbal numerical reasoning (VNR), matrix pattern recognition (Matrix) and memory pairs-matching test (Memory) as outlier tests presenting significant deviations from the expectations under the common pathway model (Figure 4). Specifically, VNR and Matrix showed stronger positive genetic associations with educational attainment and cognitive performance than those expected from the common pathway model. In contrast, the genetic correlations between these cognitive traits and ADHD were significantly more negative than those implied by the common pathway model. Follow-up models for ADHD (QTrait(4)= 31.47, *p*<.001; lSRMR=.07) and educational attainment (QTrait(4)= 40.23, *p*<.001; lSRMR=.06) with unconstrained direct paths to VNR, Matrix, and for cognitive performance (cognitive performance (QTrait(5)= 49.18, *p*<.001; lSRMR=.13) with unconstrained direct path to VNR, did not reveal significant heterogeneity, while the associations with the genetic *g* factor remained statistically significant (**Table 2**).

**Figure 4.**
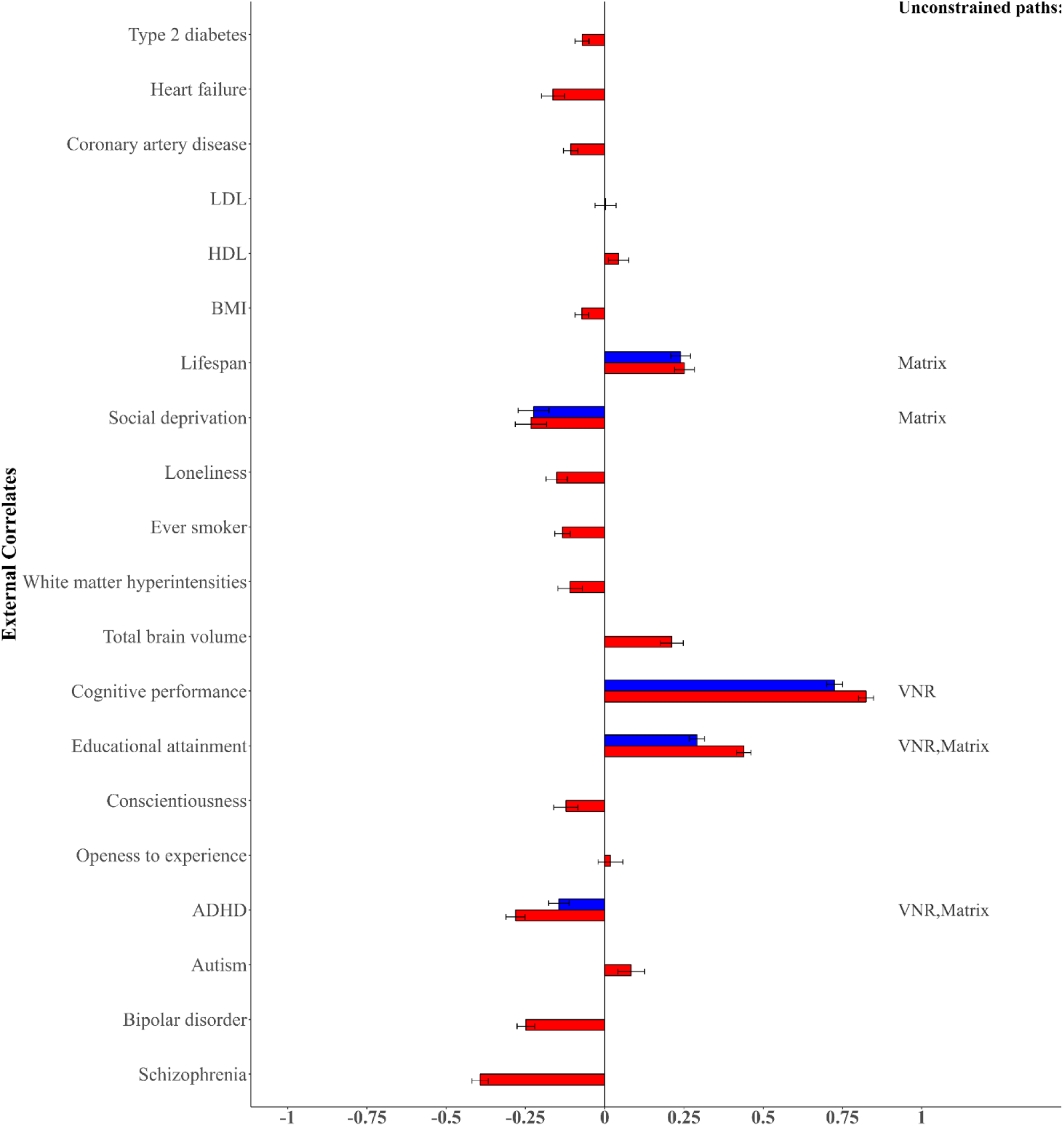
Genetic correlations across external correlates and the genetic *g* factor. The common pathway model assumes that all genetic associations between the external GWAS correlates and the individual indicator phenotypes are mediated through the common factor. The follow-up model corresponds with a model with unconstrained direct pathways to the outlier traits presenting significant deviations from the expectations under the common pathway model according to the three-test criteria of heterogeneity (see Methods).

The common pathway model estimates the genetic association between educational attainment and general cognitive function (genetic *g*) but fails to fully capture the pattern of associations across all cognitive indicators. As shown in Figure 3, both the Verbal Numerical Reasoning (VNR) and Matrix reasoning tests deviate notably from the line representing the model-implied association, exhibiting substantial residual genetic correlations with educational attainment (0.37 and 0.29, respectively). If the common pathway model adequately accounted for the genetic correlations between educational attainment and these cognitive traits, the observed correlations would scale proportionally with the indicators’ factor loadings on genetic *g*. However, VNR and Matrix reasoning display stronger-than-expected associations, suggesting the presence of additional trait-specific genetic effects. When the model is relaxed to allow for direct genetic associations between educational attainment and these two indicators, the model-implied association between educational attainment and the general cognitive factor is attenuated, decreasing from 0.439 to 0.291. These findings underscore the limitations of the common pathway model in this context and suggest that a more nuanced or multifactorial representation of genetic influences on cognitive traits may be warranted.

## Discussion

This paper extends the Genomic SEM framework by extending the QTrait method for investigating heterogeneity of genetic associations between external correlates and the indicator traits of a common factor. By incorporating both global statistical significance and effect size measures, our approach overcomes the limitations of sole reliance on statistical significance, which can either obscure meaningful differences in low-power studies or detect trivial differences in high-power studies. Furthermore, our method enables the identification of specific outlier GWAS phenotypes that deviate from the expectations of the common pathway model, revealing genetic pathways that may not be captured by shared genetic factors. This is particularly critical, as such heterogeneity could pose a threat to the external validity of genetic factor models—an issue that has often been underappreciated in the literature. By providing a more refined approach to assess how well these models approximate the genetic relationships between complex traits and external correlates, our method offers a more comprehensive and robust framework for understanding the genetic architecture of complex traits.

Combining direct case-control genome-wide association studies (GWAS) with proxy-phenotype GWAS (GWAX), such as maternal and paternal AD history, is a widely used strategy to boost statistical power in Alzheimer’s disease (AD) research (Bellenguez et al., 2022; Hujoel et al., 2020; Liu et al., 2017). However, genetic associations between AD and external biobehavioral traits, such as educational attainment (EA) and cognitive performance, can differ between direct case-control GWAS and proxy GWAX. Wu et al. (2024) employed GWAS-by-subtraction (Demange et al., 2021) to disentangle these differences, a method particularly suited for comparing pairs of traits. In contrast, our approach extends beyond pairwise comparisons, allowing for the integration of multiple indicators while automating the identification of misfit sources. Additionally, it provides a framework for testing the validity of inference within the factor model—an aspect not addressed by GWAS-by-subtraction. We observed significant and substantial heterogeneity in the genetic associations between both EA and cognitive performance, and AD. Maternal AD history displayed anomalous positive genetic correlations with both EA and cognitive performance, despite negative genetic correlations between case-control AD, cognitive performance, and EA. Our results indicate that an analysis naïve to this heterogeneity underestimated the associations between both educational attainment and cognitive performance and lower AD risk. However, this bias was automatically detected and corrected for using the function introduced here. Several hypotheses may explain divergent results with respect to direct and proxy GWAS data (de la Fuente et al., 2021; Escott-Price, V. & Hardy, J., 2022). First, proxy GWAX data could be influenced by biases in family health history reporting, such as individuals misremembering or confusing their parents’ disease status, potentially attenuating or contaminating the heritability of the proxy phenotype with other heritable traits.

Second, differences in diagnostic quality and criteria between proxy reports of historical disease status and the carefully screened case-control samples used in direct GWAS could also bias the proxy data. Finally, proxy GWAX may capture additional genetic signals not directly related to AD, including the pernicious effects of sample bias, which may itself be differential according to unmeasured heritable phenotypes (Pirastu et al., 2021; Schoeler et al., 2022). These findings and hypotheses underscore the need for careful interpretation of proxy GWAS data, distinguishing genetic influences specific to AD from those associated with other familial traits.

Our analysis of the genetic *g* factor revealed significant heterogeneity in its associations with various external correlates, shedding light on the complex genetic underpinnings of general cognitive ability. Among the 20 external correlates examined, 15 showed significant associations with the genetic g factor, and 5 of these exhibited heterogeneity, suggesting more specific genetic pathways. These included educational attainment (EA), cognitive performance, loneliness, lifespan, and attention-deficit/hyperactivity disorder (ADHD). This heterogeneity indicates that while these traits are linked to cognitive ability, their genetic influences likely vary across different cognitive domains. For example, verbal numerical reasoning (VNR) showed stronger genetic associations with educational attainment and cognitive performance than those expected by the common pathway model. This may be due to the significant crystallized knowledge component of the VNR test (Fawns-Ritchie & Deary, 2020; de la Fuente et al., 2021; Londoño et al., 2025), which aligns more closely with educational measures like math ability and degree level. Importantly, these association patterns were further clarified through outlier detection and correction. Specifically, the association between educational attainment and the genetic *g* factor was smaller in the corrected model, in contrast to the AD case, where correction led to a stronger association. This highlights that outlier detection does not systematically inflate or deflate associations, but rather corrects for bias in either direction.

While our method provides valuable insights, several limitations should be considered. First, the use of GWAS summary statistics means that our results are dependent on the quality of the underlying data, which can be influenced by factors such as sample size and bias, genotyping and phenotyping errors, and population stratification (Wainschtein et al., 2022). These factors can introduce biases, especially in studies with smaller samples or less accurate genetic data (Dastani et al., 2017). Another consideration is assortative mating, where genetic similarities between partners can inflate heritability estimates and alter patterns of genetic associations across traits, potentially distorting the observed relationships between traits (Barban et al., 2016; Border et al., 2022). These limitations highlight the importance of interpreting genetic correlations with caution and recognizing the influence of biases that may shape these associations.

In conclusion, our study introduces a novel and efficient framework for assessing the external validity of genetic factor models by evaluating how well they approximate the genetic relationships across constellations of complex traits. By incorporating both statistical significance and effect size measures, our method provides a more comprehensive understanding of genetic correlations, overcoming the limitations of traditional approaches that rely solely on statistical significance. This framework’s ability to identify genetic heterogeneity and detect outlier phenotypes not only offers valuable insights into the genetic architecture of complex traits but also facilitates the generation of mechanistic hypotheses about genetic associations. By highlighting areas where shared genetic factor models may not fully capture the underlying genetic relationships, the identification of outliers can inform targeted investigations into specific genetic mechanisms.

## Methods

### Overview of Genomic SEM

Genomic SEM is a two-stage structural equation modeling approach. In the initial stage, a genetic covariance matrix (**S**) and its associated sampling covariance matrix (**V**) are estimated using a multivariate version of Linkage Disequilibrium Score Regression (LDSC). **S** consists of heritabilities on the diagonal and genetic covariances (coheritabilities) on the off-diagonal. **V** comprises squared standard errors of **S** on the diagonal and sampling covariances on the off-diagonal, which capture dependencies between estimating errors, particularly in cases where there is participant sample overlap across GWAS phenotypes. In the second stage, a structural equation model is fitted to **S** by optimizing a fit function that minimizes the difference between the model-implied genetic covariance matrix (∑(θ)) and **S**, while considering the weights from **V**. The QTrait function employs the diagonally weighted least-squares fit function for the genomic SEM models, as described in Grotzinger et al. (2019). Validation studies conducted by Grotzinger et al. (2019) demonstrated that genomic SEM produces unbiased standard errors, appropriately accounts for sample overlap in multivariate GWAS, and provides accurate point estimates for various population-generating models.

### Overview of the QTrait function

The QTrait function is a genetically informed method for assessing the heterogeneity of associations between the genetic components of external correlates and the individual indicators of common factors representing shared genetic architecture. It includes an omnibus inferential test of heterogeneity (i.e., the QTrait statistic), which provides statistical significance, along with an effect size measure that indicates the magnitude of the heterogeneity (i.e., lSRMR). Additionally, the function incorporates an outlier detection method to identify specific indicator phenotypes that exhibit associations with external correlates not solely explained by the common factor. This feature allows the function to account for these more specific pathways, yielding more accurate parameter estimates for the association between the external correlate and the common factor. Implemented in the Genomic SEM R package (Grotzinger et al., 2019), the QTrait function uses GWAS summary data from samples of varying and unknown degrees of overlap, and does not require raw genotype data.

The required input for the function is the output from the multivariate LDSC function included in the Genomic SEM package.

The QTrait function starts by fitting a Genomic SEM common pathway model to estimate the regression coefficient relating the external correlate with the common factor (Figure 1). The function determines the statistical significance of this parameter estimate using a Bonferroni-corrected *p*- value threshold based on the number of external correlates fed to the function. If the *p*-value of the regression weight of the external correlate on the common factor exceeds the Bonferroni-corrected threshold, the function then evaluates potential heterogeneity in the associations between the external correlate and the individual indicators loading on the common factor. To do so, the QTrait function incorporates two omnibus tests of heterogeneity: the QTrait statistic, which is based on statistical significance, and the local standardized root mean squared residual (lSRMR), an effect-size measure of heterogeneity. Both indices quantify heterogeneity in the associations between the external correlate of interest and the indicator phenotypes conforming the common factor of shared genetic architecture. By integrating both statistical significance and effect size measures, the function avoids identifying negligible differences in high-powered comparisons and distinguishes meaningful differences from sampling variation in lower-powered comparisons.

### QTrait statistic: an omnibus inferential test of heterogeneity

The QTrait statistic is based on the approach by Grotzinger et al. (2022) and evaluates whether a given factor adequately accounts for the patterns of associations between an external correlate and the individual traits loading on the factor. The QTrait statistic indexes the extent of model misfit for a common pathway model in which the effects of an external correlate on the individual phenotypes are specified to occur exclusively via a single effect of the external correlate on the latent factor, compared to a less restrictive independent pathways model where effects occur directly on each phenotype. A low QTrait indicates that the external correlate plausibly influences the latent factor, while a high QTrait suggests it does not.

Under the common pathway model, the expected effects of the external correlate 𝑏_𝑐𝑜𝑟𝑟𝑒𝑙𝑎𝑡𝑒,𝐹_ on phenotype k can be expressed as 𝑏_𝑐𝑜𝑟𝑟𝑒𝑙𝑎𝑡𝑒,𝐹_×𝜆_𝑘_, i.e.,

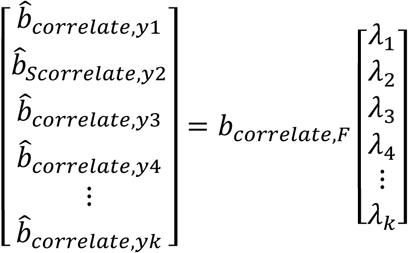

Misfit occurs when the vector of expected effects on phenotypes deviates from the vector of observed effects obtained from univariate GWAS. If the external correlate affects the individual phenotypes solely through the common factor, the vector of observed effects should be proportional to the vector of unstandardized loadings of those phenotypes, resulting in a low QTrait. Conversely, if the observed effects are not proportional to the loadings, QTrait will be high, indicating model rejection.

When QTrait is high for a given external correlate, the linear association between the vector of univariate regression coefficients and the vector of unstandardized factor loadings will be weaker, potentially indicating one or more outliers. It is important to note that we do not expect an external correlate acting exclusively on the common factor to have equal univariate associations with all phenotypes; this would only occur if the factor loadings were similar. Instead, if the external correlate truly acts exclusively on the common factor, we anticipate that the univariate associations will scale with the unstandardized factor loadings of the corresponding phenotypes. For example, a phenotype with a relatively low unstandardized factor loading should exhibit a correspondingly lower association with the external correlate compared to other phenotypes. If the association with that phenotype is comparable to associations with other phenotypes, this would contribute to a high QTrait.

A χ² difference test is used to compare the common and independent pathways models, as indicated by the QTrait heterogeneity index, which follows a χ² distribution with k−1 degrees of freedom (where k represents the number of traits loading on the common factor). A significant QTrait index suggests that the common factor does not adequately explain the pattern of associations between the individual traits and the external correlate. We note that we explore the regression coefficients of external correlates instead of Z statistics or *p*-values when investigating heterogeneity because differences in sample sizes, heritabilities, or polygenicity can lead to Z statistics that do not correlate with unstandardized factor loadings.

### Local standardized root mean squared residual correlation as an effect size measure of heterogeneity

The local standardized root mean squared residual (lSRMR) is an effect-size-based measure of heterogeneity, adapting the local measure of parameter difference across groups (localSRMD) introduce by Schwaba et al. (2023) for comparing differences in Genomic SEM parameters across subgroups of individuals (e.g. males and females). Unlike the original localSRMD, which focuses on parameter differences, the lSRMR computes the root mean squared residual correlation. This metric acts as a local index of heterogeneity, comparing the observed genetic correlations—derived from the LDSC output—with the correlations implied by the common pathway model:

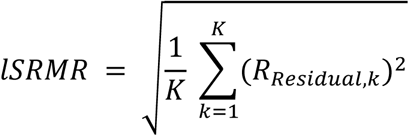

Where K is the number of indicator traits that load on the common factor, and RResidual,k corresponds with the residual genetic correlation for indicator *k*, calculated as

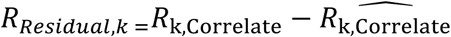

Where 𝑅_k,Correlate_ represents the observed LDSC-derived genetic correlation between the external correlate and the k-th indicator, and 𝑅_k,_^Ĉ^_orrelate_ is the model-implied genetic correlation between the external correlate and the same indicator.

The lSRMR serves as a global standardized effect size index of heterogeneity, requiring two thresholds for meaningful interpretation. To detect significant heterogeneity, the lSRMR must exceed both a context-specific threshold and an absolute threshold (default = 0.10). The default context-specific threshold is set as 25% of the root mean square genetic correlation between the external correlate and the individual indicator phenotypes that load on the common factor, ensuring that heterogeneity is evaluated relative to the observed effect sizes. This threshold strikes a balance between sensitivity and robustness, identifying non-trivial heterogeneity while minimizing noise. In addition, the absolute threshold provides a fixed benchmark to ensure that only substantial deviations are considered meaningful, regardless of the dataset’s specific context. Both thresholds must be surpassed to indicate meaningful heterogeneity in the genetic correlations, suggesting that the observed associations between the traits and the external correlate cannot be fully explained by the model.

### Outlier detection and follow-up models

The QTrait function incorporates a method for identifying outliers that deviate from the expectations under the common pathway model. These outliers correspond to traits for which the common factor does not adequately capture the association pattern with the external correlate.

The outlier detection method identifies outliers based on the following two criteria related to residual genetic correlations: 1) the residual genetic correlation for indicator k (i.e., Rresidual,k) exceeds a predefined absolute threshold (the default value is 0.10), and 2) Rresidual,k exceeds a predetermined proportion of the root mean square genetic correlation between the external correlate and the indicator phenotypes loading on the factor (the default value is 0.25, as per the lSRMR). Specifically, a trait k is considered an outlier if it satisfies the following conditions:

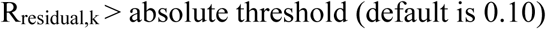

and

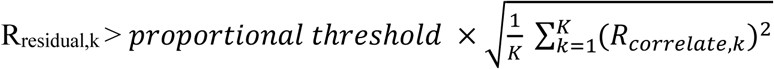

Outlier detection follows the same approach as the global effect-size-based heterogeneity assessment provided by the lSRMR, requiring two thresholds for meaningful identification. The lSRMR and outlier detection both rely on the context-specific threshold and the absolute threshold. For outlier detection, the default context-specific proportional threshold is set as 25% of the root mean square genetic correlation, ensuring that outliers are evaluated relative to observed effect sizes, balancing sensitivity and robustness. The absolute threshold ensures that only substantial deviations are considered meaningful, avoiding the introduction of unnecessary model parameters for minor fluctuations. Both thresholds must be exceeded to identify meaningful outlier traits, ensuring that direct paths are added only when the evidence justifies such model modifications. This dual approach maintains model parsimony while effectively detecting significant outliers in a context-sensitive manner, adaptable across different datasets.

When significant heterogeneity is detected via the three-criteria test, the QTrait function proceeds to fit a follow-up model that includes an unconstrained direct path from the external correlate to the most extreme outlier. If the genetic correlation between the external correlate and the common factor remains significant in this follow-up model, and significant heterogeneity persists, the function selects the most extreme outlier identified in that model and repeats the process until either the number of direct paths reaches saturation (i.e., k-1 direct paths are estimated, where k is the number of traits comprising the factor) or no significant heterogeneity is observed.

### Sources of GWAS summary data for empirical analyses

#### Cognitive Function

We drew GWAS summary statistics obtained from the UK Biobank for seven cognitive tests: trail-making tests-B (n = 78,547), tower rearranging (n = 11,263), verbal numerical reasoning (VNR, n = 171,304), symbol digit substitution (n = 87,741), memory pairs-matching test (n = 331,679), matrix pattern recognition (n = 11,356), and reaction time (RT, n = 330,024). We employ the same common factor model described by de la Fuente et al. (2021), where a general dimension of shared genetic liability, termed genetic *g*, was identified in these same cognitive traits. Detailed information on the specific tests, participant criteria, and assessment details can be found in de la Fuente et al. (2021).

#### Alzheimer’s Disease

We compiled summary data from three published European ancestry direct case-control GWAS of AD and GWAX of parental history of AD. The direct case-control GWAS summary statistics included the discovery sample from the IGAP consortium (Kunkle et al., 2019), comprising 21,982 Clinical AD cases (mean age of onset = 72.93 years) and 41,944 controls (mean age of evaluation = 72.415 years). To address potential bias arising from variations in case prevalence across these cohorts, we adopted the approach outlined by Grotzinger et al. (2022), using the sum of effective sample sizes (4vk(1−vk))nk, where vk is the sample prevalence for each contributing GWAS to the meta-analysis). The GWAX summary data for the proxy-phenotype AD included 27,696 cases and 260,980 controls for the maternal history of AD and 14,338 cases and 245,941 controls for the paternal history of AD, with both phenotypes sourced from the UK Biobank (Marioni et al., 2019). Case status in the UK Biobank was determined by responses to questions about parental Alzheimer’s disease/dementia history during the initial assessment visit (2006–2010), the first repeat assessment visit (2012–2013), and the imaging visit (2014+). Participants with parents younger than 60 years, those who died before reaching 60 years, and those lacking parental age information were excluded. Zhang et al. (2020) reported mean ages of 83.7 years for maternal cases, 81.8 years for paternal cases, 78.1 years for maternal controls, and 76.2 years for paternal controls. Further details on case ascertainment, genotyping, and quality control can be found in the original articles providing the corresponding summary statistics.

### External correlates

To explore the generality versus specificity of associations between the general intelligence and the Alzheimer’s disease factors, we selected external GWAS correlates based on their relevance for cognitive function, health related lifestyle and general liability for disease. The external correlates included educational attainment (n = 3,037,499; Okbay et al., 2022), cognitive performance (Nn 257,841; Lee et al., 2018), Total Brain Volume (n = 36,778; Fürtjes et al. 2023), White Matter Hyperintensities (n = 50,970; Sargurupremraj et al., 2020), Ever smoker (n= 518,633; Liu et al., 2019), Loneliness (n= 452,302 ; Day et al., 2018), Social deprivation (n = 112,151; Hill et al., 2016), parental lifespan (n = 1,012,240; Timmers et al., 2019), BMI (n= 681,275;Yengo et al., 2018), HDL (n = 1,320,016; Graham et al., 2021), LDL (n = 1,320,016; Graham et al., 2021), Coronary artery disease (n = 1,378,170;Van der Harst & Verweij, 2018), Heart failure (n = 933,970; Shah et al., 2020), and Type 2 diabetes (n = 933,970; Mahajan et al. 2022).

For the specific analysis of the genetic *g* factor, we included neuropsychiatric and personality traits given their relevance for cognitive function reported in the literature. Neuropsychiatric disorders data were obtained mostly from the Psychiatric Genetics Consortium (PGC) database (Sullivan, 2010), including schizophrenia (Neff = 117,494; Trubetskoy et al., 2022), bipolar disorder (Neff = 101,963; Mullins et al., 2021), autism spectrum disorder (Neff = 43,778; Grove et al., 2019), attention deficit hyperactivity disorder (Neff = 103,135; Demontis et al., 2023). Personality traits were also considered. The GWAS summary statistics for Openness to experience (n = 220,015) and Conscientiousness (n = 234,880) were taken from the Million Veteran Program (MVP) (Gaziano et al., 2016) release version 4 as described in Gupta et al. (2024). These traits were measured using a 10-item scale (BFI-10) as part of a self-report Lifestyle survey provided to MVP participants. Only data from individuals of European ancestry were used, with a mean age of approximately 65.5 years for each trait, and 8% of the sample was female.

